# An *in-vivo* digital atlas of the spatially reliable cerebral vasculature in mice

**DOI:** 10.1101/2021.10.21.465264

**Authors:** Rukun Hinz, Meriam Malekzadeh, Lore M. Peeters, Verdi Vanreusel, Marleen Verhoye, Annemie Van der Linden, Georgios A. Keliris

**Author notes:** **Corresponding author** Prof. Georgios Keliris, Bio-imaging lab, University of Antwerp, Campus Drie Eiken– Building Uc 1.07, Universiteitsplein 1 -2610 Wilrijk – Belgium.

## Abstract

Cerebral vascular abnormalities can have a large impact on brain function and have been frequently detected as a comorbidity in various neuropathologies. The mouse is the most common pre-clinical animal model used to investigate neuropathologies and thus cerebral vascular atlases of this species are indispensable. In particular, an atlas derived from multiple animals and that can address inter-subject variability is still absent. The current study presents a mouse cerebral vascular atlas developed from MR 2D-TOF angiograms acquired from a relatively high number (N=21) of 12 weeks old male C57BL/6 mice. The vascular atlas we present depicts large arteries, veins and sinuses that are more spatially reliable across animals than smaller vessels. The atlas is available for download and contains multiple datasets: i) a cerebral vascular atlas; ii) a digital segmentation of large vessels (arteries and veins); iii) a high resolution T2 anatomical background aligned to the atlas; iv) a skeletonized vascular atlas; and v) a vascular graph model. We suggest that these components provide a very potent analysis tool for mouse fMRI data and that the atlas can be used as a template for multimodal brain imaging studies. To this end, we also demonstrate an explicit application of the atlas for the investigation of vascular influences in resting-state fMRI.

## 1. Introduction

The cerebral vasculature has an indispensable supporting role in maintaining the energy homeostasis of the brain by supplying it with oxygen and nutrients as well as clearing the brain of waste metabolites. [1, 2]. Any structural aberrations in the cerebral vasculature, such as those due to stroke, can deplete the continuous blood supply and can result in cell death and loss of function. Furthermore, structural vascular abnormalities have been shown to commonly arise as comorbidities in various neuropathological disorders, such as Alzheimer’s disease, Huntington’s disease and multiple sclerosis [3–5]. Hence, appropriate sensitive tools to investigate cerebral vasculature are required.

In preclinical research, mice models are by far the most commonly used animal models to study neuropathological disorders. During the last decades, technological advancements have been made in preclinical imaging allowing for the characterization of the mouse brain’s vasculature at different spatial scales. For example, *ex-vivo* imaging techniques like knife-edge scanning microscopy (KESM), micro-optical sectioning tomography (MOST), serial two photon tomography and light sheet microscopic (LSM) imaging have been used to acquire high-resolution imaging data of the vascular architecture up to the capillary level [6– 11]. These techniques result in the acquisition of large datasets that allow in-depth analysis of the whole brain vasculature. Furthermore, *in-vivo* imaging techniques, such as computed tomography (CT), magnetic resonance angiography (MRA) and transcranial functional ultrasound (fUS) have been optimized to visualize cerebral vasculature with its main advantage that it can follow-up vascular integrity in the same subject over time [12–14]. These latter techniques predominantly image the macro vasculature, but CT and MR imaging can also be performed *ex-vivo* with longer acquisitions and higher resolution to acquire information about the microvasculature [15, 16]. Using the aforementioned techniques, several vascular atlases have been developed to assist in identifying the mouse brain vasculature. In one study, *ex-vivo* CT imaging was combined with MRI to image the mouse whole brain vasculature of four mice together with an anatomical background [17]. This atlas provides the segmentation of the vasculature at the macroscopic level allowing the identification of most large vessels. A more detailed vascular atlas was obtained in the study of Xiong et al. using micro-optical sectioning tomography (MOST) [18]. In that study, the entire mouse brain vascular architecture was mapped of five animals and up to the capillary level allowing the identification of the whole brain vasculature as well as the investigation of microvascular densities in the brain.

Knowledge of the vascular architecture is not only important for identifying vascular alterations in disease states but can also play a key role in supporting a variety of neuroimaging techniques. Imaging studies such as fUS and intrinsic optical imaging (IOS) are exclusively based on acquiring vascular information, but with limited spatial resolution [19, 20]. Vascular atlases that would be suitable to use across studies and incorporate anatomical information can help to identify the spatial location of the vessel. Furthermore, vessel occlusion studies that investigate stroke can benefit from vascular atlases to select the appropriate vessel to occlude or to estimate the size of the hypoxic area. An important number of studies that can benefit from vascular atlases are those using functional MRI (fMRI) and/or resting state fMRI (rsfMRI). These techniques rely on the acquisition of blood oxygenation level dependent (BOLD) signals that are mainly derived from the hemodynamic responses caused by the neurovascular coupling [21]. However, studies have shown that other vascular effects, such as vascular drainage, also influence the BOLD signal [22, 23]. Additionally, resting state signals have been shown to be confounded by signals from large arteries and veins [24]. These confounding signals are thought to be systemically produced from outside the brain and seem to be propagating throughout the blood vessels [25]. A vascular atlas that can be readily used in such studies, can help in identifying potential vascular confounds and assist in further analysis steps aiming to remove them.

To be able to apply a vascular atlas in the aforementioned studies, it is assumed that the spatial location of the vasculature is conserved between subjects. However, the study of Xiong et al. has shown that the spatial location of small vessels (< 40 µm) can be highly variable between subjects of the same strain, while larger vessels (> 40 µm) are more spatially reliable [18]. An efficient method to visualize spatially reliable vessels is to develop a vascular atlas based on angiograms from multiple subjects. As such, spatial unreliable vessels will be suppressed by averaging the datasets. However, mouse vascular atlases to date focused on the visualization of the detailed vasculature in a single mouse or a small number of subjects and were created *ex-vivo* in order to use techniques that allow high resolution [18, 26]. Thus, existing mouse vascular atlases are less suitable for application across studies.

In this study, we developed and present an *in-vivo* acquired cerebral vascular atlas of the large vessels using 21 male C57BL/6 mice of 12 weeks of age. To this end, non-invasive imaging, 2D time of flight (TOF) MRA was used and combined with high-resolution anatomical T2-weighted MRI for spatial identification. Our digital atlas is provided together with an anatomical MRI template, and thus allows easy and direct co-registration to MRI datasets or can be used as a reference template for segmenting newly acquired individual angiography. We also provide a skeletonized version of the atlas that can be used as a graph to calculate parameters such as path length. Moreover, in this study, we demonstrate an explicit application of the atlas for the investigation of vascular influences in resting-state fMRI by estimating the overlap between vasculature and resting state networks (RSNs).

## 2. Material and Methods

### 2.1. Animals and ethical statement

The study was conducted using 12 weeks old male C57BL/6 wild-type mice (N=21) (Janvier). Mice were kept under a day/night cycle of 12h/12h respectively, with an average room temperature of 20-24°C and 40% humidity. All mice were group housed with *ad libitum* access to water and standard rodent chow. All applicable institutional and/or national guidelines for the care and use of animals were followed. Procedures were performed in accordance with the European Directive 2010/63/EU on the protection of animals used for scientific purposes. The protocols were approved by the Committee on Animal Care and Use at the University of Antwerp, Belgium (permit number: 2016-10) and all efforts were made to minimize animal suffering. Experiments have been reported in compliance with the ARRIVE guidelines.

### 2.2. Data availability statement

Data used in this study are available online and can be downloaded at www.uantwerpen.be/en/research-groups/bio-imaging-lab/research/mri-atlases/. Datasets include 2D-TOF MRA of each individual subject, vascular atlas, skeletonized version of the vascular atlas and the anatomical study-specific template.

### 2.3. Imaging procedure

All MR imaging procedures were performed on a 9.4 T Biospec MR system (Bruker, Germany) with a receive-only 4 channel array cryoprobe (Bruker, Germany) combined with a volume transmit resonator. All animals underwent MRA imaging, whereas a partial group (N=13) was scanned a second time to extract rsfMRI data.

#### 2.3.1. MRA imaging

For MRA imaging, animals were first anesthetized using 5% isoflurane in a gas mixture of 30% O2 and 70% N2 for induction, which was then subsequently decreased to 2% isoflurane for maintenance during the remainder of the session. All mice were placed head-fixed in the scanner by using ear- and tooth-bars and ophthalmic ointment was applied to the eyes to prevent desiccation. During the experimental procedure, animals’ physiology was closely monitored. As such, respiratory rate was followed using a pressure sensitive pad (MR-compatible Small Animal Monitoring and Gating system, SA Instruments, Inc.). Body temperature was followed using a rectal thermistor and was maintained at (37.0 ± 0.1) °C using a feedback controlled warm air heat system (MR-compatible Small Animal Heating System, SA Instruments, Inc.). Additionally, blood oxygenation was monitored using a pulse oxygenation meter (MR-compatible Small Animal Monitoring and Gating system, SA Instruments, Inc.).

MRI data acquisition consisted out of the measurement of three orthogonal oriented 2D-TOF-MRA to assess the macrovasculature with blood flow in each direction: (FOV: (20×20) mm^2^, matrix dimension (MD: [320×320], 110, 130, 90 slices respectively in the coronal, axial and sagittal oriented scans, slice thickness (ST): 0,3 mm, slice shift: 0.15 mm, voxel dimension (0.0625×0.0625×0.15) mm^3^, TE/TR: 3.2/17 ms, flip angle: 60°,number of averages: 3). Subsequently, a high-resolution 3D T2-weighted Turbo-RARE was acquired for the development of an anatomical template (FOV: (20×20×15) mm^2^, MD: [256×256×192], voxel dimension = (0.078 mm)^3^, TE/TR: 40/2500 ms, RAREfactor: 16). After the imaging procedure, animals were kept in a recovery box with infrared heating until they were fully awake to compensate for post-anaesthethic decreased body temperature.

#### 2.3.2. Resting state functional MRI

RsfMRI was acquired 1 to 2 weeks after the MRA imaging session. To do so, animals’ anesthesia was first induced with 5% isoflurane in a gas mixture of 30% O_2_ and 70% N_2_ and was lowered to 2% isoflurane for maintenance. Mice were head-fixed and were monitored similarly as for the previous MRA session. Ophthalmic ointment was applied to the eyes. After fixation, animals received a subcutaneous bolus of medetomidine (0.05 mg/kg) which was 15 minutes later followed by a subcutaneous continuous infusion of medetomidine (0.1 mg/kg/h). After bolus injection, isoflurane concentration was gradually decreased to 0.4% isoflurane over a time period of 5 minutes.

In each animal, three orthogonally placed T2-weighted Turbo-RARE (FOV: (20×20) mm^2^, MD: [256×256], 12 slices, ST: 0.4 mm, TE/TR: 33/2500 ms, RAREfactor: 8) anatomical images were acquired to ensure slice position uniformity between animals. Next, a coronal oriented T2-weighted anatomical reference scan was acquired using a Turbo-RARE sequence (FOV: (24×16) mm^2^, MD: [256×256], 16 slices, ST: 0.5 mm, TE/TR: 33/2500 ms, RAREfactor: 8) to be used as an anatomical background for functional imaging data. Afterwards, B0 field maps were acquired to estimate magnetic field inhomogeneity that was used for the subsequent local shimming procedure within a rectangular volume of interest covering the mouse brain. Thirty minutes post-bolus injection, rsfMRI scans were acquired using a T2*-weighted single shot echo planar imaging (EPI) sequence (FOV: (24×16) mm^2^, MD: [96×64], 16 slices, ST: 0.5 mm, voxel dimension: (0.25 × 0.25 × 0.50) mm^3^, TE/TR: 16/1000 ms, 600 volumes). After the imaging procedure, mice received a subcutaneous bolus injection of 0.1 mg/kg atipamezole (Antisedan, Pfizer, Karlsruhe, Germany) to counteract the effects of medetomidine and then the animals were placed in a recovery box with infrared heating until they were fully awake.

### 2.4. Data Processing

#### 2.4.1. Anatomical Template

The 3D-T2 weighted anatomical scans of each subject were used to develop a study-specific anatomical template. All co-registration processing was performed using Advanced Normalization Tools (ANTS). Scans were first debiased to equalize signal intensities over the FOV. Afterwards, the template was created using the standard ANTS create template protocol.

#### 2.4.2. MRA

The 2D-TOF MRA sequence is an excellent tool for non-invasive imaging of the complete mouse brain vasculature. This sequence allows for the detection of vascular flow perpendicular towards the slice direction. Therefore, three orthogonally oriented 2D-TOF MRAs were used to visualize vascular flow in the rostral-caudal (fig.1A), dorsal-ventral (fig.1B) and left-right (fig.1C) direction. To develop subject specific angiograms, scans were first visually inspected for motion artifacts and then combined using a maximum Intensity Projection (MIP) (fig.1D). Afterwards, each subject vascular angiogram was co-registered to the anatomical template by using the previously estimated subject to template transformation parameters from the template creation step. Finally, co-registered vascular angiograms of each subject were averaged across all subjects to create a vascular atlas.

**Figure 1.**
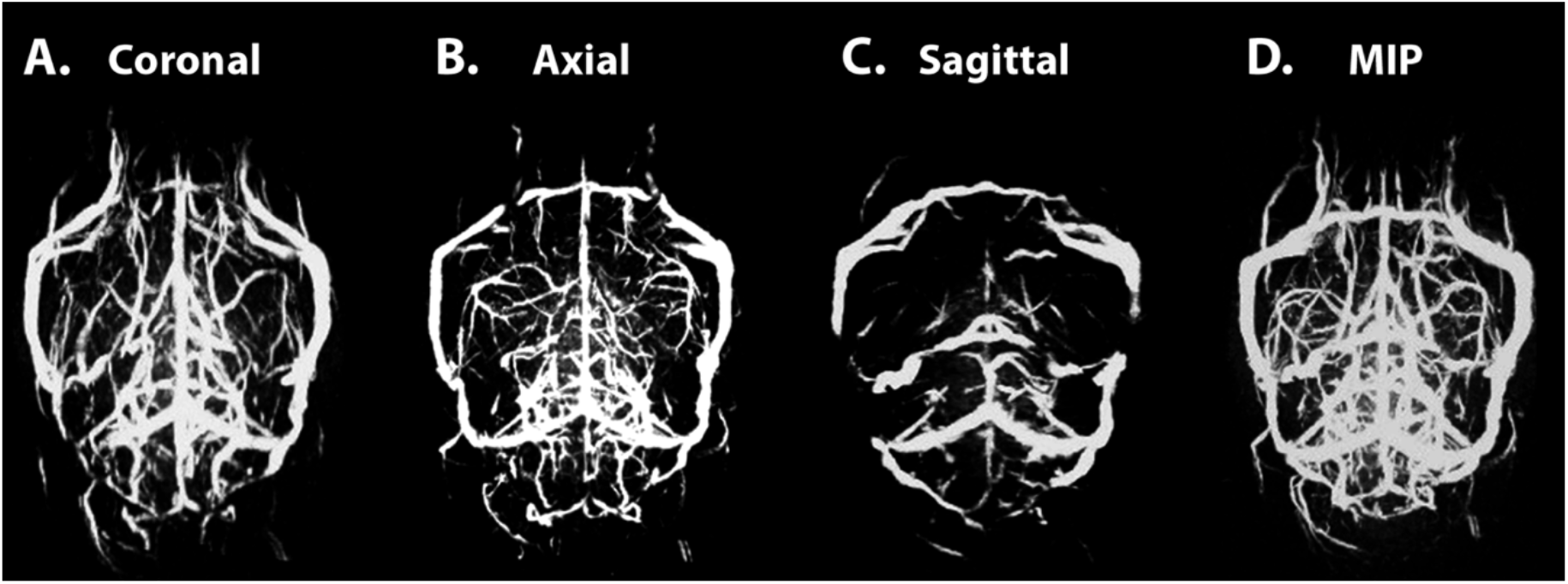
Time of flight magnetic resonance imaging. Example of a 2D-TOF MRA dataset of a representative animal. Dataset included three orthogonally positioned 2D-TOF MRA scans with the slice package placed (A) Coronally, (B) Axially and (C) Sagittally allowing the visualization of vascular flow in the rostral-caudal, dorsal-ventral and left right direction respectively. Datasets were combined using a maximum intensity projection (MIP) to visualize whole brain vasculature (D).

Next, processing of the data was performed in Amira 5.4 (FEI, Thermo Scientific). Individual blood vessels were manually delineated on the vascular atlas and labeling was based on previous reported vascular atlases [17, 18]. Afterwards, the vascular atlas was skeletonized using Amira 5.4 skeletonization tool transforming the developed atlas into a nodes and edges model. This model was then saved as centerline voxels in Amira 5.4 which converts the nodes and edges model into single voxel lines representing the inner core of the vessels. Additionally, the nodes and edges model allowed for automatic clustering of enclosed circuits. As the circuits of large arteries and veins did not overlap, the nodes and edges model facilitated the automatic separation of arteries and veins. Information coming from the nodes and edges model was then used to develop a vascular graph model in Matlab code (MATLAB R2019a, The MathWorks Inc. Natick, MA, USA).

#### 2.4.3. Resting state functional MRI

Data processing was performed using SPM 12 (Statistical Parametric Mapping, http://www.fil.ion.ucl.ac.uk), Gift toolbox (Group ICA of fMRI toolbox version 3.0a: http://icatb.sourceforge.net) and Matlab 2014a (MATLAB R2014a, The MathWorks Inc. Natick, MA, USA). Functional data was first realigned using a 6-parameter (rigid body) spatial transformation. Next, images were used to create a study-specific EPI template by normalizing the first volume EPI of each subject to the first volume of a single subject by using a global 12-parameter affine transformation followed by a non-linear transformation and an averaging of all normalized scans. Afterwards, individual subject functional datasets were normalized to this study-specific EPI template using a global 12-parameter affine transformation and subsequently a non-linear transformation. Normalized data was then smoothed in-plane by using a Gaussian kernel with a full-width half-maximum of twice the voxel size (0.5 × 0.5 mm). Data were then further band pass filtered between 0.01-0.15 Hz.

To extract RSN from rsfMRI data, an independent component analysis (ICA) was performed with the Infomax algorithm with 15 pre-defined number of components. Components representing RSN were calculated by performing a one-sample t-test to obtain distinct networks and identified by comparison to previously observed RSNs [27–30]. ICA data was then co-registered from the EPI template space to the vascular atlas high-resolution 3D-T2 weighted anatomical template space by performing a rigid, affine and non-linear transformation using the ANTS toolbox. For this transformation, each voxel index from the lower resolution EPI data was used as a value instead of a binary mask so that we could calculate overlap with the original EPI voxels after nearest neighbor interpolation. To identify RSNs with potential vascular influences, RSNs were sorted based on the percentage overlap with the vascular atlas. To this end, for each RSN, the vascular overlap fraction (VOF) was calculated by dividing the total amount of (unique) voxels overlapping with the vascular skeleton by the total amount of unique voxels of that RSN.

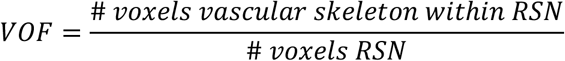

Similarly, we calculated partial VOF for specific vascular compartments (i.e. arterial, venous/sinous) or specific vessels by replacing the total vascular skeleton with the skeleton of the vascular compartment of interest

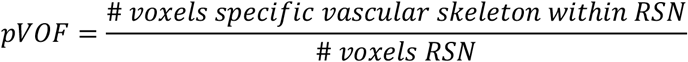

After sorting, the two RSNs with the highest vascular overlap were selected and further analyzed in order to identify the specific arteries and vessels overlapping with these RSNs. To this end, the percentage of vessel inclusion (PVI) within each network was calculated for each artery and vein by dividing the number of voxels of each RSN overlapping with each vessel’s skeleton with the number of voxels of that vessel’s skeleton and converting to percentage.

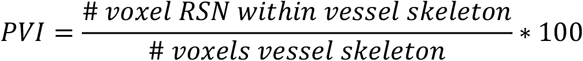

## 3. Results

### 3.1. Vascular atlas

The 2D-TOF MRA data of 21 male C57BL/6 wild-type mice was used for the development of the vascular atlas. The segmentation resulted in delineation of the main large vessels that were spatially reliable across the sample of 21 mice. Furthermore, the segmentation was sorted into arteries (fig.2) and veins/sinuses (fig.3) to have a clear description of the vascular architecture (further described below). Given the limited spatial resolution of *in-vivo* MRA as well as the inter-subject small variability in vessel positions, the vessel sizes were substantially overestimated. To overcome this limitation, centerlines of the blood vessels were extracted by performing a skeletonization on the segmented data (fig.4). These centerlines depict the vascular atlas in single voxel lines representing the inner core of the blood vessels, which is also the position of highest inter-subject spatial reliability. Given this property, such an atlas representation is especially useful for investigating vascular effects of specific vessels in imaging data. Moreover, the skeletonization also allowed the development of a vascular graph, which represents the hierarchical structure of the vasculature and allows more explicit types of analyses, such as intervascular distance between vessels (or vessels and voxels), branching order, parent-child relationships, *et cetera*.

#### 3.1.1. Arteries

The major arterial system of the mouse brain is visually depicted in figure 2A and its hierarchical graph (fig. 2B) gives an overview of its connections. The blood supply to the mouse brain is provided via four main arteries *i*.*e*. (i) the two **vertebral arteries** that mainly supply blood to the arteries of the brain stem and the cerebellum and (ii) the two **internal carotid arteries** that mainly supply blood to arteries feeding the subcortical and cortical brain regions.

**Figure 2.**
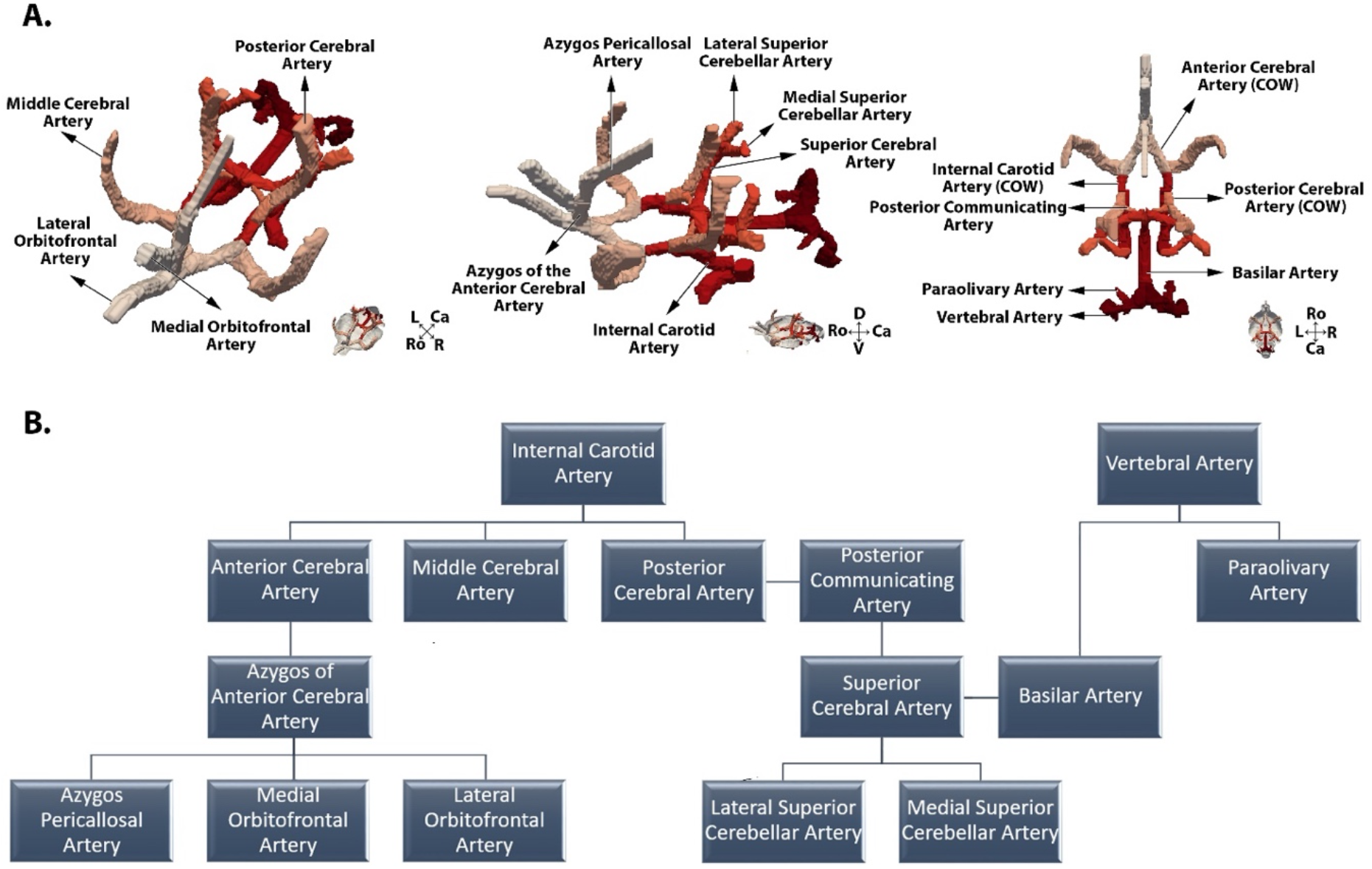
Arterial System and vascular graph. A. Three-dimensional overview of the arterial system with annotations included in the vascular atlas. Crosshair indicates rostral-caudal (Ro-Ca), dorsal-ventral (D-V) and left-right (L-R) direction. B. Hierarchical structure and annotations of the mouse brain arterial system.

The **vertebral artery** depicted one clear branch giving rise to the **para-olivary artery** providing blood towards the inferior olive region. The two vertebral arteries then merge in the midline into the **basilar artery** that is located on the ventral surface of the brainstem until it reaches the pons/midbrain junction to give rise to two **superior cerebral arteries**. These superior cerebral arteries end up in the **lateral and medial superior cerebellar artery**.

The **internal carotid artery** branches in rostral, lateral and caudal direction resulting in the **anterior, middle and posterior cerebral artery** respectively. The **anterior cerebral artery** of both internal carotid arteries heads rostrally on the ventral part of the brain and merges together medially to form the **azygos of the anterior cerebral artery**. This artery is directed dorsally and provides blood to the frontal lobe of the brain, including the olfactory bulb and orbitofrontal regions by branching into the **lateral orbitofrontal artery** and later the **medial orbitofrontal artery**. Next, the **azygos of the anterior cerebral artery** curves rostral and gives rise to the **azygos pericallosal artery** that runs over the dorsal part of the corpus callosum delivering blood to posterior-medial parts of the frontal-, parietal and temporal lobes. The **middle cerebral artery** branches off from the lateral side of the internal carotid artery and extends over the surface of the ventral part of the brain where it feeds blood to the hypothalamus, caudate putamen and amygdala. It then continues in the dorsal direction on the surface of the brain to provide blood to most of the cerebral cortex. The **posterior cerebral artery** is dorsally directed where it is providing blood towards the caudal part of the cerebrum including the hippocampus and the superior colliculus as well as cortical areas. Furthermore, the posterior cerebral artery connects with the superior cerebral artery via the posterior communicating artery.

At the ventral part of the brain around the level of the optic chiasm and the hypothalamus, the arterial system forms a circulatory anastomosis called the **circle of Willis** (COW). This anastomosis connects the internal carotid artery and the vertebral artery. The circle of Willis is built from the **basilar artery**, which connects to both **superior cerebral arteries** that are both connected with the **posterior cerebral arteries** via the **posterior communicating artery**. The two **posterior cerebral arteries** are each connected with an **internal carotid artery** that gives rise to the **medial cerebral artery** and the **anterior cerebral artery**. The anterior cerebral artery of both sides fuse into a circular anastomosis.

#### 3.1.2. Veins and Sinuses

The major mouse-brain venous system produces a drainage system to remove blood containing low nutrients as well as waste metabolites from the brain. The 2D-TOF MRA detected the dorsal venous system as well as the deep venous system. Furthermore, veins outside of the brain were observed and labeled. The venous system of the mouse brain is visualized in figure 3A and its venous graph (fig. 3B) gives an overview of the connections of each vein.

**Figure 3.**
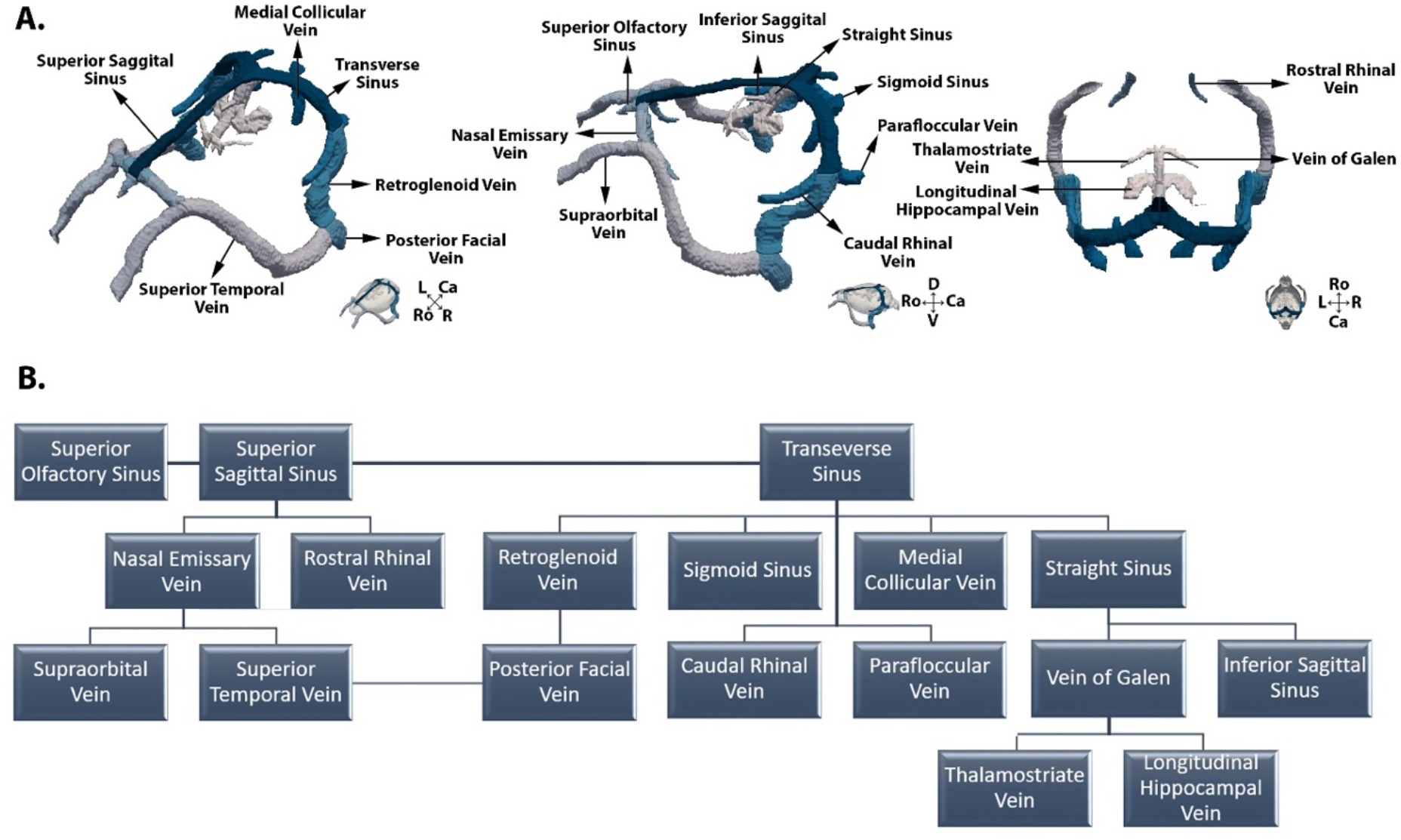
Venous System and vascular graph. A. Three-dimensional overview of the venous system with annotations included in the vascular atlas. Crosshair indicates rostral-caudal (Ro-Ca), dorsal-ventral (D-V) and left-right (L-R) direction. B. Hierarchical structure and annotations of the mouse brain venous system.

The detected dorsal venous system contains mainly four sinuses and their connecting veins. The most rostral detected sinus is the (i) **superior olfactory sinus** which is connected in the caudal direction with the (ii) **superior sagittal sinus**. These sinuses are located dorsally on the pial surface and cover the midline of the cerebral cortex brain. In the rostral end, the superior sagittal sinus produces the **rostral rhinal veins** in the ventral direction that reaches the frontal lobe. At the caudal end, the superior sagittal sinus splits into two lateral-ventral oriented (iii) **transverse sinuses** that are each connected with a caudal medial oriented (iv) **sigmoid sinus**. The aforementioned sinuses cover most of the pial surface of the cerebral cortex. The transverse sinus was observed to have three branches *i*.*e*. a ventral directed **medial collicular vein** that penetrates into the cerebral tissue to reach the colliculus areas, the **parafloccular vein** that is directed caudally and reaches the paraflocculus, and finally the rostral directed **caudal rhinal vein**.

The deep venous system branches off from the medial-dorsal part of the transverse sinus and gives rise to the **straight sinus**. The sinus is rostroventrally directed and gives rise to the **inferior sagittal sinus** as well as the **vein of Galen**. The vein of Galen was then observed to produce two lateral oriented **longitudinal hippocampal veins** and further caudally two **thalamostriatal veins**.

The venous blood is removed from the brain via venous outflow systems. Here the 2D-TOF MRA detected the dorsal and caudal outflow. The dorsal outflow starts from the **nasal emissary vein**, which is connected with the rostral part of the superior sagittal sinus. This vein is oriented laterally and merges together with the **supraorbital vein** coming from the eyes into the **superior temporal vein**. The caudal outflow system occurs from the **retroglenoid vein** that is connected to the transverse sinus. The retroglenoid vein and the superior temporal vein merges into the **posterior facial vein** that will eventually remove the venous blood from the head via the external jugular vein.

#### 3.1.3. Skeletonization

The developed 2D-TOF MRA atlas overestimates the vessel size. This is especially clear when comparing the anatomical background images, where vessels produce hypo-intense signals, with the 2D-TOF MRA data (fig. 4B). To overcome this, the vascular atlas was further processed to only visualize the centerlines of each vessel using a skeletonization process. By doing so, the whole brain vasculature could be represented by nodes and edges model or with center lines with a width of one voxel (fig.4). Furthermore, this skeletonization process allows for the development of a vascular graph model (see fig. 2B, 3B), visualizing the hierarchical structure of the vascular architecture.

**Figure 4.**
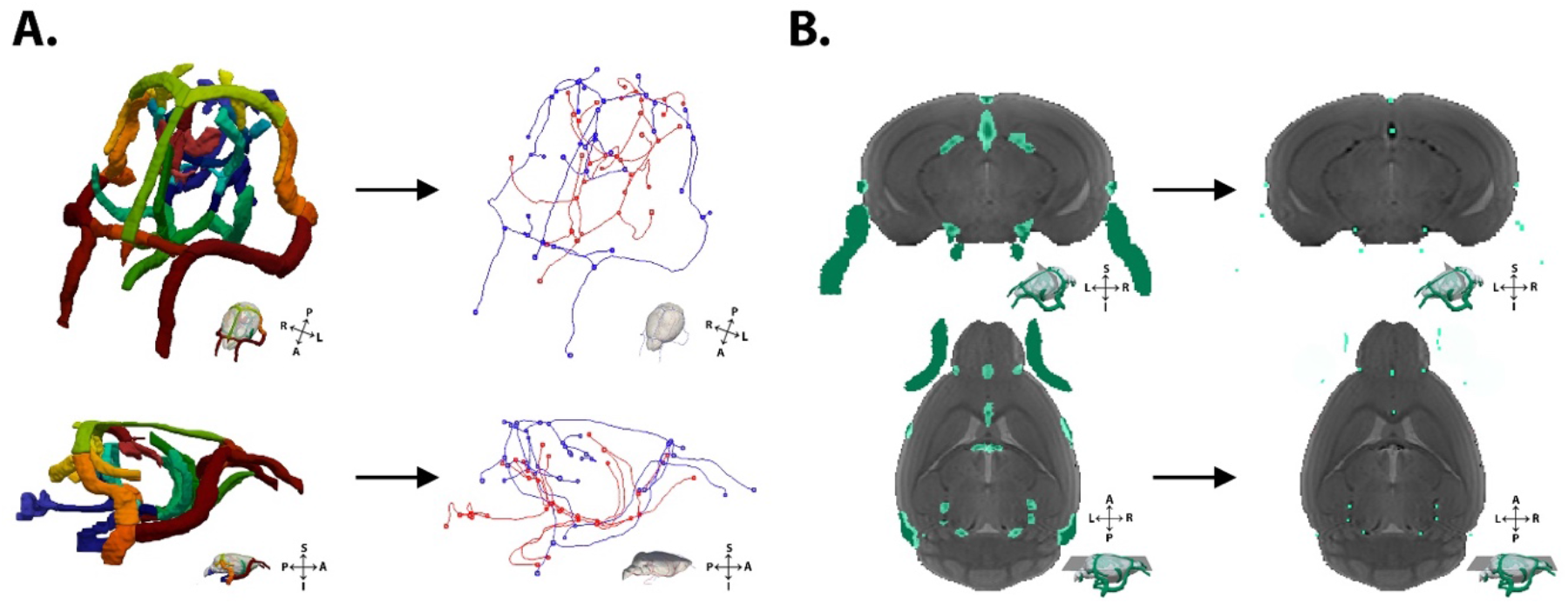
Skeletonization of the mouse brain vasculature. A. The labeling of the vasculature was skeletonized and is represented in nodes and edges. The nodes represent starting, end or bifurcation points, while edges indicate vessel paths. Furthermore, the skeletonization process allows for fast detection of arteries (red) and veins (blue) as both were detected as separate circuits B. When comparing the labeled vasculature with the anatomical template, a discrepancy between vessel size could be observed. Centerlines can be used to demonstrate the presence of a vessel irrespectively of its size. For visualization purposes centerlines were depicted with a centerline thickness of 4 voxels.

### 3.2. Resting state functional connectivity

Functional MRI sequences that rely upon the detection of the BOLD contrast such as gradient echo EPI, are known to suffer from vascular influences especially due to large vessels. In this study, rsfMRI data from a subset of animals used to build the reported atlas, were used to investigate the extent of overlap between RSNs and the major mouse brain vasculature. The RSNs were extracted from functional MRI data using independent component analysis (ICA) and revealed nine functionally relevant networks (suppl. fig. 1). To calculate the percentage overlap between the vasculature and the identified RSN components, we used the skeletonized version of our atlas. Given that the skeletonized atlas has a single voxel resolution, the overlap essentially finds voxels of the resting state network in which major blood vessels are present. The results indicated that a number of networks demonstrated substantial fraction of major vascular overlap (suppl. table 1). Specifically, the VOF values were as follows sorted from highest to lowest overlap: Cingulate-Retrosplenial I = 0.06, Cingulate – Retrosplenial II = 0.0483, Orbitofrontal = 0.0428, Hippocampal = 0.0417, Striatum = 0.024, Somatosensory barrel field = 0.0239, Somatosensory – Motor = 0.0184, Primary somatosensory = 0.0183, Visual Cortex = 0.0167. The two networks with the highest VOF were both networks which include the cingulate and retrosplenial cortex *i*.*e*. cingulate-retrosplenial (Cg-Rs) I and II (fig. 5). Specifically, Cg-Rs I and Cg-Rs II components demonstrated values of around 6% and 5% of voxels with overlap with major mouse vasculature respectively. The Cg-Rs I network was observed to include the complete cingulate cortex and partially the retrosplenial cortex as well as motor cortex, preoptic area, dorsal thalamus, superior colliculus and hippocampus. The Cg-Rs II network contained only the posterior part of the cingulate, the entire retrosplenial cortex as well as the parietal association and visual cortex. The substantial overlap of these networks with the major mouse vasculature can directly be appreciated on the 3D visual representation of these components as shown in figure 6 (fig. 6). Another two networks demonstrated substantial vascular overlap of over 4% of voxels was the orbitofrontal and hippocampal networks, while all other networks demonstrated vascular overlap values < 2.5%.

**Figure 5.**
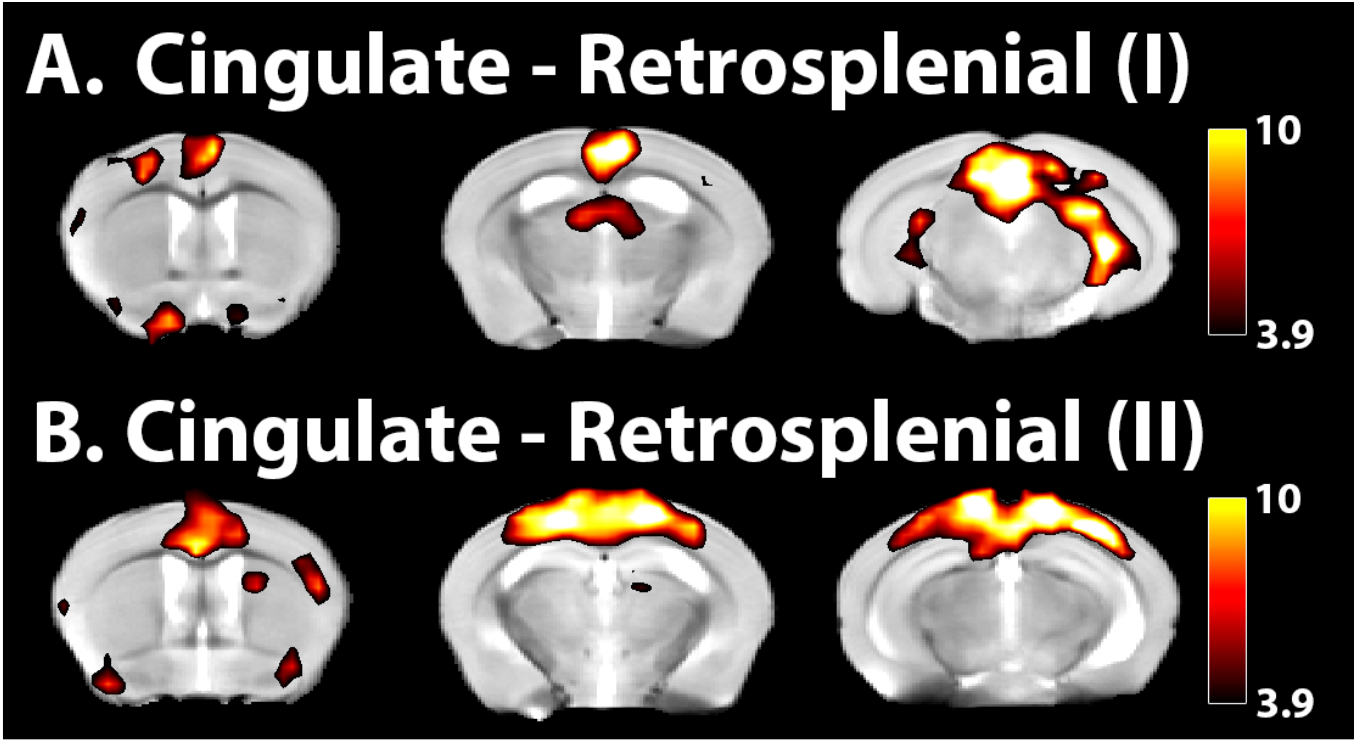
Resting state networks with highest vascular overlap. Statistical map of two resting state networks with highest vascular overlap (A) Cingulate-Retrosplenial I (0.21%) and (B)II (0.17%). Statistical maps were produced by one sample t-test (p<0.001, uncorrected) performed on the outcome of an independent component analysis. Color bar represents t-values.

**Figure 6.**
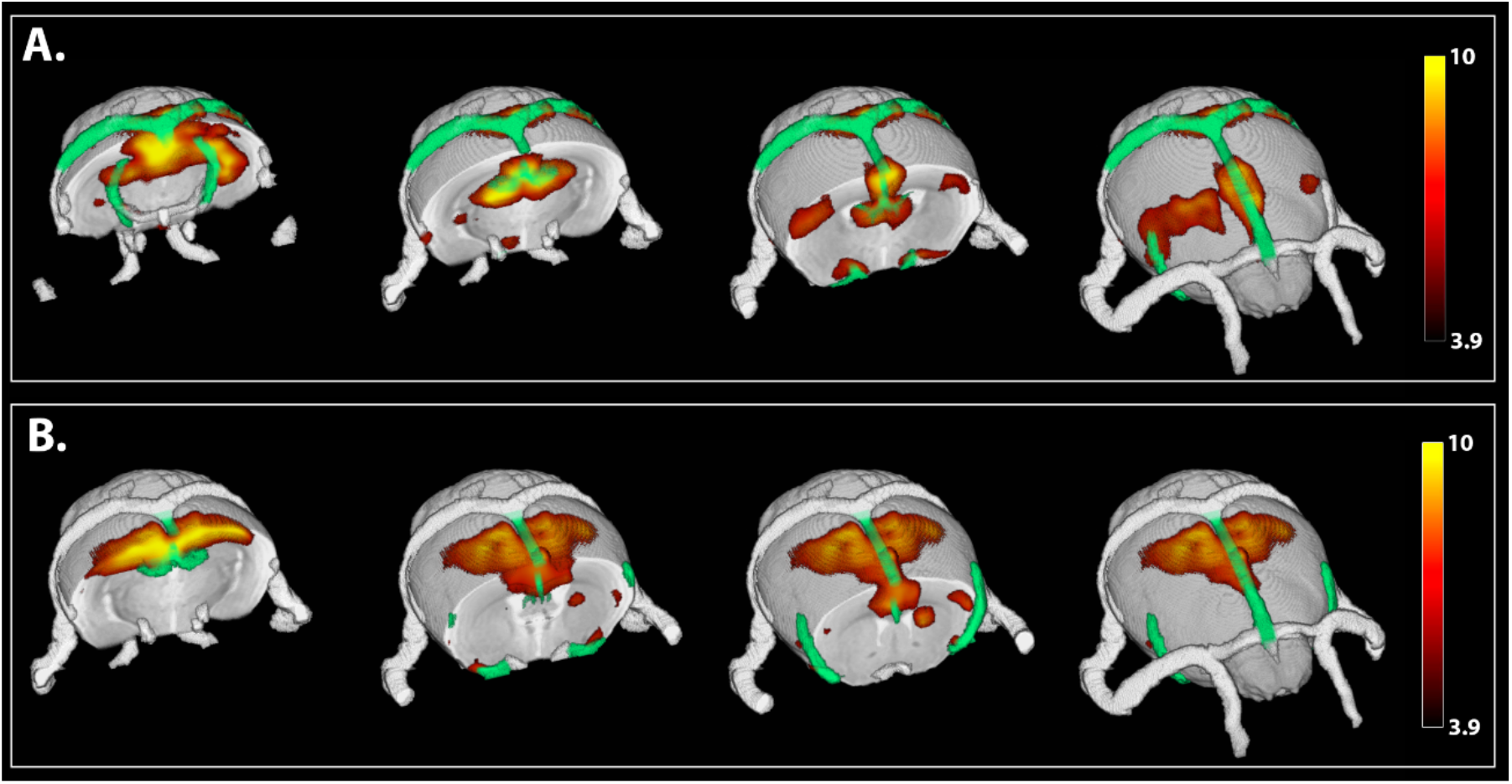
Resting state networks and vascular overlap. 3D visual representation of the statistical maps of (A) cingulate-retrosplenial I and (B) II network together with the brain vasculature (MIP). Statistical maps were produced by one sample t-test (p<0.001, uncorrected) performed on the outcome of an independent component analysis. Color bar represents t-values. Blood vessel that partially overlap with the resting state network were highlighted in green.

To better understand the contribution of the arterial and venous/sinous compartments as well as the contribution of specific vessels within each of the two cingulate – retrosplenial networks, we also calculated the partial VOF (pVOF) for each vessel (table 1).

**Table 1.** Percent Vascular Inclusion (PVI) in Cingulate-Retrosplenial I and II. Percentage vessel of the vascular skeleton contained within the resting state component.

| Vessel Name | Cingulate - Retrosplenial<br>I | Cingulate - Retrosplenial<br>II |
| --- | --- | --- |
|  | (% vessel within<br>component) | (% vessel within<br>component) |
| Internal Carotid Artery (Right) | 26.67 | / |
| Internal Carotid Artery (Left) | 28.57 | / |
| Superior Cerebral Artery (Right) | 15.38 | 5.13 |
| Lateral Superior Cerebellar Artery (Right) | 78.57 | / |
| Lateral Superior Cerebellar Artery (Left) | 52.38 | / |
| Posterior Communicating Artery (Right) | 8.33 | / |
| Posterior Cerebral Artery (Right) | 25.00 | / |
| Posterior Cerebral Artery (Left) | 21.57 | / |
| Middle Cerebral Artery (Right) | 11.59 | 10.14 |
| Middle Cerebral Artery (Left) | 5.63 | 5.63 |
| Anterior Cerebral Artery (Right) | 64.29 | / |
| Anterior Cerebral Artery (Left) | 31.03 | / |
| Azygos of Anterior Cerebral Artery | 27.27 | / |
| Azygos Pericallosal Artery | 41.82 | 50.91 |
| Medial Orbitofrontal Artery | 14.29 | 2.86 |
| Superior Sagittal Sinus | 13.58 | 24.53 |
| Transverse Sinus (Right) | / | / |
| Transverse Sinus (Left) | 22.73 | / |
| Medial Collicular Vein (Left) | 7.41 | / |
| Straight Sinus | 100.00 | 92.31 |
| Inferior Sagittal Sinus | 100.00 | 100.00 |
| Vein of Galen | 79.17 | 100.00 |
| Longitudinal Hippocampal Vein (Right) | 100.00 | 100.00 |
| Longitudinal Hippocampal Vein (Left) | 100.00 | 41.67 |
| Thalamostriate Vein_(Right) | 50.00 | 3.85 |
| Thalamostriate vein (Left) | 50.00 | 10.00 |

The Cg-Rs I network includes arteries, veins and sinuses while Cg-Rs II contained mainly veins and sinuses. In the Cg-Rs I, a venous overlap could be seen from the superior sagittal, transverse and the straight sinuses including all its labeled branches. The arterial overlap of the Cg-Rs I involved the internal carotid artery, anterior cerebral artery and its branches, the posterior/middle cerebral artery and the superior cerebellar artery. The Cg-Rs II vascular overlap contained mainly venules *i*.*e*. the superior sagittal sinus and the straight sinus and its branches. Arterial overlap was limited to the superior/middle cerebral artery, azygos pericallosal artery and the medial orbitofrontal artery.

## 4. Discussion

An intact vascular architecture is essential for the healthy function of the brain. It is therefore of utmost importance for many studies to be able to evaluate the contribution of vessel signals and/or detect vascular abnormalities. Previously developed mouse cerebral vascular atlases mainly focused on the visualization of the - as much as possible - complete vascular tree down to the small vessels and capillaries. This resulted in high-resolution large datasets that tend to be time consuming to acquire and difficult to process [31] and therefore report results from a limited number of animals [17, 18]. In the current study, we developed of a mouse cerebral vascular atlas that focusses on visualizing the spatially reliable larger vessels in a high number of C57BL/6 wild-type mice (N=21) at the age of 12 weeks. In contrast to previous atlases, this atlas was specifically developed to be used as a reference vascular atlas for group analysis across studies that use C57BL/6 wild-type mice. By using high number of animals, the averaging of the individual cerebral vasculature enhances spatial reliable vessels and suppresses vessels demonstrate anatomical variations across subjects. In addition, the vascular atlas is co-registered to high-resolution anatomical data which is adding spatial information to the vasculature.

### 4.1. 2D-TOF MRA

The vascular atlas was developed using a 2D-TOF MRA sequence. This MRI sequence was specifically chosen for its high sensitivity towards the vasculature and its complete non-invasiveness, which allows for easy implementation in future MRI studies. The method allowed for the detection of blood vessels based upon their flow perpendicular to the slice position. It was therefore required to acquire three orthogonally positioned 2D-TOF MRA scans to obtain the complete overview of the vasculature. The 2D-TOF MRA was chosen instead of its 3D-TOF MRA counterpart due to its superior sensitivity to slow flow allowing for the detection of a wider range of vessels. Moreover, 3D-TOF MRA is also more sensitive to motion artifacts that could compromise the data acquisition. By combining three 2D-TOF MRA datasets, we implemented a method for non-invasive reproducible detection of *in-vivo* cerebral vasculature in mice.

### 4.2. Vascular atlas

The presented *in-vivo* vascular atlas contains multiple datasets: i) a cerebral vascular atlas; ii) a digital segmentation of large vessels (arteries and veins/sinuses); iii) a high resolution T2 anatomical background; iv) a skeletonized vascular atlas; and v) a vascular graph model.

The cerebral vascular atlas is an average vascular template created by all co-registered 2D-TOF MRA datasets and is accompanied by a T2-weighted anatomical template. These datasets can be used as a co-registration endpoint to acquire either vascular information or anatomical information. The templates will be especially useful for studies that acquire 3D anatomical information (MRI, CT) and want to extract cerebral vascular information or for studies that rely on the acquisition of vascular information (fUS, IOS) and want to get anatomical information. The vascular atlas was manually segmented, which resulted in the labeling of spatially reliable vessels. The segmentation included the main large arteries, veins and sinuses within the brain as well as their outflow systems going around the head. However, small branches were not included due to spatial variation between subjects. The anatomical structure of the vessels was similar as in the paper of Xiong et al. that produced a high resolution *ex-vivo* vascular atlas in the same C57BL/6 mouse strain [18]. Small differences were found between our labeling and the vascular atlas of Dorr et al. that mainly revolved around the posterior communicating artery being connected with the posterior cerebral and internal carotid artery, while in our dataset it was connecting the posterior and superior cerebral artery [17]. However, this can be explained as the aforementioned atlas was acquired in a different mouse strain *i*.*e*. CBA mouse strain. Previous studies have reported structural cerebral vascular differences between mouse strains [32, 34, 35]. This indicates caution when using the vascular atlas in other strains than the C57BL/6 strain. Furthermore, due to the spatial variability of the cerebral vasculature, the segmentation of the blood vessels were overestimated. To overcome this, the segmentation was skeletonized and vascular centerlines were extracted. As the vascular centerlines depicts blood vessels as single voxel lines it is a more appropriate tool to investigate the spatial location of the vasculature. The centerlines will therefore be of use in studies that are particularly interested in defining the presence of vasculature in their datasets. In addition to the centerlines, the skeletonization process results in a representation of the vasculature as nodes and edges allowing for the development of a vascular graph model. The vascular graph model contains the hierarchical structure of the cerebral vascular architecture and allowed automatic clustering of the arterial and venous system. This model can be used to get insights in blood flow patterns which could be valuable for studies performing BOLD and perfusion analysis.

### 4.3. Application of the vascular atlas in rsfMRI

RsfMRI relies upon the detection of fluctuations in the BOLD signal, which is assumed to be an indirect measurement of changes in neural activity detected via the neurovascular coupling. As BOLD signals are based on detecting changes in blood oxygenation, it is assumed that it is most sensitive to the capillaries where most of the oxygenation exchange occurs. However, the BOLD signal has been shown to be influenced by the macrovasculature [22]. More specifically, confounding BOLD signals of vascular origin have been widely detected coming from downstream draining veins [22]. This is caused due to the accumulation of the blood coming from the capillaries to the draining veins and thus smearing out the BOLD signal via the venules. Therefore, as an application of the vasculature atlas, we implemented the atlas to rsfMRI data to investigate the vascular contribution in the observed RSN. The vascular overlap analysis could show that the two highest overlapping components were components covering connections between the cingulate and retrosplenial cortex (Cg-Rs I and II). These regions are commonly part of the default mode-like network in mice [27, 29, 36]. Using the vascular atlas, we could reveal that the Cg-Rs I & II RSNs have the strongest vascular contribution, but this is still limited to around 6% and 5% of the original resolution EPI voxels being penetrated by the large, reliable vessels included in our atlas. This indicates that even if these networks suffer from slightly higher vascular influences in comparison to some other networks, the majority of their voxels are not penetrated by the large vessels and thus their functional correlations most probably reflect neuronal components.

## 5. Conclusion

In conclusion, this article presents a newly developed *in-vivo* mouse brain vascular atlas based on 21 adult male C57BL/6 wild-type mice (12 weeks old) acquired with MRI. The atlas includes: i) a cerebral vascular atlas; ii) segmentation of large vessels; iii) a high resolution T2 anatomical background; iv) a skeletonized vascular atlas; and v) a vascular graph model that are available online for download. The atlas acts as a co-registration template for studies that require both vascular and anatomical information. As a potential application, we superimposed the atlas on rsfMRI data of adult male C57BL/6 wild-type mice which enabled us to identify a differential degree of overlap between the vasculature and the different RSNs. Undoubtedly the use of this atlas will contribute to a more solid interpretation and post processing of resting state data obtained in mice.

## 6. Acknowledgments

This research was supported by the fund of Scientific Research Flanders (grant agreement G048917N), Flagship ERA-NET (FLAG-ERA) FUSIMICE (grant agreement G.0D7651N),, Molecular Imaging of Brain Pathophysiology (BRAINPATH) under Grant Agreement Number 612360 within the Marie Curie Actions-Industry-Academia Partnerships and Pathways (IAPP) program. MRI equipment was funded by the Flemish Impulse funding for heavy scientific equipment (granted to AVdL). The computational resources and services used in this work were provided by the HPC core facility CalcUA of the Universiteit Antwerpen, the VSC (Flemish Supercomputer Center), funded by the Hercules Foundation and the Flemish Government – department EWI.

## Author contribution statement

Rukun Hinz: Study initiation, Data acquisition, Data processing, Writing of the article

Meriam Malekzadeh: Data processing: vascular segmentation

Lore M. Peeters: Study initiation, Data processing, Writing of the article

Verdi Vanreusel: Data processing: Co-registration

Marleen Verhoye: Study initiation, Writing of the article

Annemie Van der Linden: Study initiation, Writing of the article

Georgios A. Keliris: Study initiation, Data Processing, Writing of the article, Study supervision

