## Supplementary information for "An *in-vivo* digital atlas of the spatially reliable cerebral vasculature in mice"

***Corresponding author**

Prof. Georgios Keliris

Bio-imaging lab, University of Antwerp

Campus Drie Eiken– Building Uc 1.07

Universiteitsplein 1 -2610 Wilrijk – Belgium

**Keywords:** Digital cerebral vascular atlas, Mouse, Resting-state functional MRI, Vascular skeleton, Magnetic Resonance Imaging, Angiography, TOF MRA

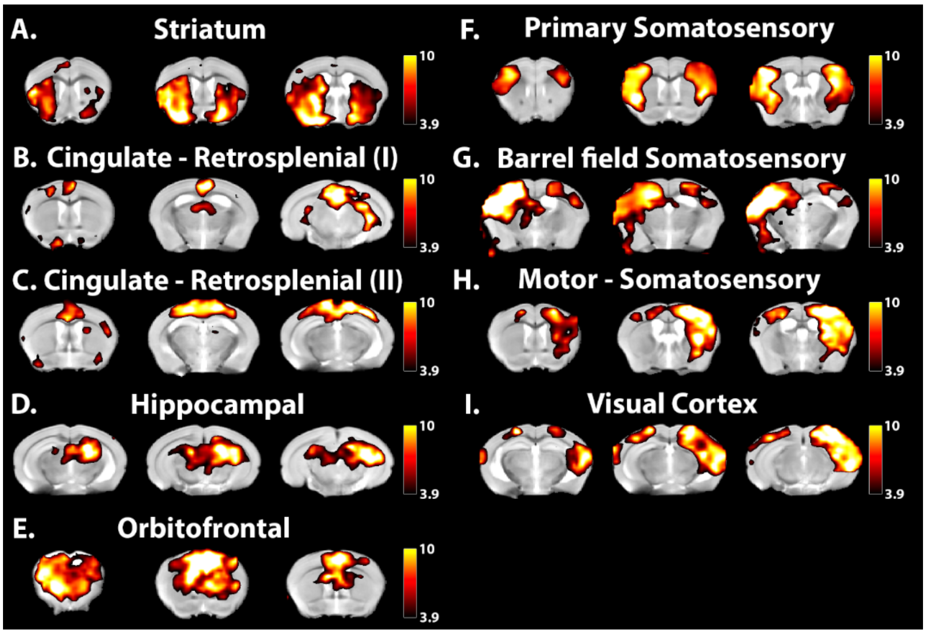


**Supplementary figure 1. ICA detected resting state networks.** *Statistical map of all detected resting state networks. Statistical maps were produced by one sample t-test (p<0.001, uncorrected) performed on the outcome of an independent component analysis. Color bars represent T statistic values.*

| ICA Component | Vascular Overlap Fraction (VOF) |
|---|---|
| **Cingulate - Retrosplenial I** | 0.0600 |
| **Cingulate - Retrosplenial II** | 0.0483 |
| **Orbitofrontal** | 0.0428 |
| **Hippocampal** | 0.0417 |
| **Striatum** | 0.0240 |
| **Barrel field Somatosensory** | 0.0239 |
| **Motor - Somatosensory** | 0.0184 |
| **Primary Somatosensory** | 0.0183 |
| **Visual Cortex** | 0.0167 |

**Supplementary Table 1. Overlap of vascular skeleton with resting state components.** *Vascular Overlap Fraction (VOF) of vascular skeleton with RSN. Values were calculated by dividing the number of voxels where vascular skeleton overlaps with the RSN by the total amount of voxels in that RSN.*
